# Effects of advanced age upon astrocyte-specific responses to acute traumatic brain injury in mice

**DOI:** 10.1101/845883

**Authors:** Alexandria N. Early, Amy A. Gorman, Linda J. Van Eldik, Adam D. Bachstetter, Josh M. Morganti

## Abstract

**Background:** Older-age individuals are at the highest risk for disability from a traumatic brain injury (TBI). Astrocytes are the most numerous glia in the brain, necessary for brain function, yet there is little known about unique responses of astrocytes in the aged-brain following TBI.

**Methods:** Our approach examined astrocytes in young adult, 4-month-old, versus aged, 18-month-old mice, at 1, 3, and 7 days post-TBI. We selected these time points to span the critical period in the transition from acute injury to presumably irreversible tissue damage and disability. Two approaches were used to define the astrocyte contribution to TBI by age interaction: 1) tissue histology and morphological phenotyping, and 2) transcriptomics on enriched astrocytes from the injured brain.

**Results:** Aging was found to have a profound effect on the TBI-induced loss of homeostatic astrocyte function needed for maintaining water transport and edema – namely, aquaporin-4. The loss of homoeostatic responses was coupled with a progressive exacerbation of astrogliosis in the aged brain as a function of time after injury. Moreover, clasmatodendrosis, an underrecognized astrogliopathy, was found to be significantly increased in the aged brain, but not in the young brain. As a function of TBI, we observed a transitory refraction in the number of these astrocytes, which rebounded by 7 days post-injury in the aged brain. The transcriptomics found disproportionate changes in genes attributed to reactive astrocytes, inflammatory response, complement pathway, and synaptic support in aged mice following TBI compared to young mice. Additionally, our data highlight that TBI did not evoke a clear alignment with previously defined “A1/A2” dichotomy of reactive astrogliosis.

**Conclusions:** Overall, our findings point toward a progressive phenotype of aged astrocytes following TBI that we hypothesize to be maladaptive, shedding new insights into potentially modifiable astrocyte-specific mechanisms that may underlie increased fragility of the aged brain to trauma.

## Background

Traumatic brain injury (TBI) is one of the most powerful environmental risk factors for the subsequent development of Alzheimer’s disease (AD) and related dementias [1–7]. Despite the aged population being at the greatest risk for acquiring a TBI, as well as the significant and long-lasting neurologic sequelae following the initial trauma in the aged population, current knowledge regarding how the aged brain responds to TBI remains disproportionately low, especially with respect to cell-specific responses. In the current study, we focused on the response of astrocytes to TBI in the aged brain to begin to elucidate several cell-specific dynamics. It is well known that astrocytes play a critical role in maintaining homeostasis in the CNS; tightly regulating ATP, glucose and glutamate, synaptic pruning and function, while also serving a crucial role for blood brain barrier (BBB) integrity and cerebral blood flow (CBF) [8–15]. However, as a response to injury or disease, astrocytes are able to rapidly respond, initially via ATP-mediated calcium signaling, in a generalized description referred as reactive astrogliosis [16–18]. In the young-adult brain, contemporary evidence has shown that astrocytes play a critical neuroprotective role following TBI, as chemically ablating the vast majority of GFAP^+^ astrocytes has been shown to exacerbate neuronal loss and perpetuate inflammatory response [19], owing to several intrinsic functions of astrocytes in mediating damage repair following TBI.

A continuing growth of supporting evidences implicate astrocytes as active participants in the multicellular networked responses potentially underlying, resolving, or exacerbating CNS diseases and injury [20, 21]. As a variety of roles of astrocytes within the disease or injury milieu are being identified, it is becoming clearer that astrocyte response is rather heterogenous [17, 20, 22–24], with implications pointing toward a diversity of graded responses [25]. Critically, among these responses are multiple domains encompassing inflammatory response, tissue protection, vascular response, as well as neuronal functionality [26]. However, despite the recognition of the diverse actions of astrocytes in response to trauma, relatively little information exists on how normal aging may alter the capacity of astrocytes to mount an appropriate response to TBI. Recent evidence has shown, as a consequence of normal aging, the brain acquires a chronically elevated inflammatory phenotype, which has been shown to negatively impact astrocyte function, leading toward their increased inflammatory bias [27–29]. Aged astrocytes were also shown to disproportionately respond to an inflammatory challenge (e.g. LPS), compared to young [28]. Collectively, these works proposed that the aged brain promotes a “neurotoxic” bias in astrocytes predominated by a pro-inflammatory response, with permissive phenotypes supporting increased synaptic clearance and neuronal damage [28, 29]. In parallel, we have recently demonstrated that the aged brain promotes exacerbated neuroimmune responses following TBI [30, 31]. However, the understanding of the complex intrinsic and extrinsic mechanisms governing astrocyte-specific responses to TBI remains limited, especially in the aged brain.

Ultimately, examining TBI in aged animal models are critically necessary to begin to unravel both intrinsic and extrinsic cell-specific responses to TBI, in order to more accurately understand the pathophysiological mechanisms likely to be found in the population at greatest risk for acquiring a TBI.

## Methods

### Animals

All experiments were conducted in accordance with the National Institutes of Health *Guide for the Care and Use of Laboratory Animals* and were approved by the Institutional Animal Care and Use Committee of the University of Kentucky. Young adult (3-month-old; Jackson Laboratory) and aged adult (17-month-old; NIA aged rodent colony) male and female *C57BL6* mice were used for all experiments. Prior to experimental procedures, all mice were acclimated to housing conditions at the University of Kentucky for approximately 4 weeks. All mice were group housed 4-5 per cage in individually ventilated cages, in environmentally controlled conditions with standard light cycle (14:10hr light:dark cycle at 21°C) and provided food and water *ad libitum*. Animals’ ages at the time of surgery were approximately 4 or 18 months.

### Surgical procedure

All animals were randomly assigned and divided as equally as possible between sexes to their treatment group. Animals were anesthetized with 2.5% isoflurane before having their scalp shaved. Mice were maintained with 2.5% isoflurane via a non-rebreathing nose cone coupled to a passive exhaust system connected to a stereotaxic surgical frame (Stoelting). Animals’ heads were secured to the stereotaxic frame using Delrin non-traumatic ear bars (Stoelting), eye ointment was applied, and scalps disinfected using betadine solution. Animal temperature during surgical procedures was maintained by a heating pad set to 37°C. A mid-line incision was made through the scalp to expose the skull. All mice received a craniectomy approximately 3.5mm in diameter using a microburr electric drill with the center point to the coordinates of −2.0mm (anteroposterior), 2.0mm (mediolateral), with respect to bregma. The controlled cortical impact (CCI) injury was reproduced using the Leica electromagnetic impactor and a 3.0mm convex tip, as we have previously described [30, 32, 33]. Impact parameters were as follows; impact velocity of 4.0m/s, dwell time of 0.3s, to a depth of −0.9mm, rotated 20° on the vertical axis to match the curvature of the brain. Following surgery, scalps were closed using surgical staples and mice were transferred to a recovery cage placed on top of a heating pad. Animals were monitored until they were fully ambulatory, as exhibited by the resumption of movement and grooming behaviors. Surgical procedures were conducted in batches, such that mice were killed for all endpoints at approximately the same time during each day. Sham surgical controls were killed to match CCI post-surgical intervals, therefore shams represent a mix of 1, 3, and 7-day intervals. All animals fully recovered from surgical procedures.

### Histological tissue collection and preparation

At the predefined post-injury interval, mice were anesthetized with 5.0% isoflurane before exsanguination and transcardial perfusion with ice-cold phosphate buffered saline (PBS), followed by 4% paraformaldehyde in PBS. Immediately following perfusion, whole brain tissues were removed and bisected along the midline, and the ipsilateral hemisphere was drop-fixed in 4% buffered paraformaldehyde (PFA) overnight at 4°C. Following the overnight post fixation, brain hemispheres were transferred into 30% sucrose for at least 4 days before sectioning on a microtome to a 30μm thickness. Tissue sections were serially collected at a 1:10 interval into 2mL screw cap tubes containing 30% sucrose and stored at - 20°C.

### Astrocyte immunofluorescent labeling

One 2mL tube comprising every tenth serial section through the ipsilateral hemisphere was stained per animal using the following procedures. Sections were washed at room temperature three times for 10 minutes each in PBS, followed by 1 hour of blocking using 10% normal goat serum (NGS) in PBS containing 0.1% TritonX-100 (PBST). Subsequently, sections were incubated with primary antibodies against GFAP (Thermo Fisher Scientific Cat# 13-0300, RRID:AB_2532994, 1:800 dilution), S100B (Agilent Cat# GA50461-2, RRID:AB_2811056, 1:400 dilution), Vimentin (Millipore Cat# AB5733, RRID:AB_11212377, 1:400 dilution), or aquaporin-4 (Aqp4) (Sigma-Aldrich Cat# HPA014784, RRID:AB_1844967, 1:400 dilution) diluted in 1% NGS-PBST. Primary incubations were conducted overnight, approximately 16 hours, at 4°C. Sections were subsequently washed at room temperature 5 times in PBST for 5 minutes each. Using appropriate detection antibodies, diluted in 1% NGS-PBST, for rabbit (Thermo Fisher Scientific Cat# A-11034, RRID:AB_2576217), rat (Thermo Fisher Scientific Cat# A-21094, RRID:AB_2535749), or chicken (Thermo Fisher Scientific Cat# A-11041, RRID:AB_2534098) each at 1:200 dilutions, sections were incubated for 2 hours at room temperature. Tissue sections were subsequently washed 3 times in PBST for 5 minutes each, followed by 2 washes in PBS for 5 minutes each. Sections were mounted on Superfrost slides (Fisher #12-550-15) and allowed to air dry in the dark overnight before coverslipping with Antifade mounting medium with DAPI (Vector #H1200). Slides were sealed with nail polish and allowed to dry overnight.

### Cellular imaging and quantification

Immunofluorescently labeled sections were imaged using a Zeiss Axio Scan Z1 digital slide scanner at 20x magnification. Digital images of each slide and its sections were analyzed for threshold-defined pixel-positive area fraction using Halo Image analysis software (Indica Labs, v2.3.2089.34) utilizing the AreaQuantification v1.0 algorithm. For all analyses, an investigator blinded to the study conditions outlined anatomical region consisting of the dorsal hippocampus. The positive staining baseline was thresholded against representative young-sham positive staining, such that any pixel at that intensity or greater (e.g. brighter) was quantified. The number of positive pixels was normalized per area outlined for each section to account for outlined region-to-region area variability. All sections were batch analyzed using the stored parameters in the algorithm. For quantification of clasmatodendrosis, dorsal hippocampal structures were outlined as above, however a blinded investigator used Halo Annotation tool to manual mark each cell. The number of annotations was normalized per area for each tissue section. Additionally, representative confocal images of clasmatodendrosis in astrocytes were acquired using a Nikon C2Plus Confocal Microscope. Representative renderings of the GFAP, Vimentin, and S100B positive surfaces were created using Imaris (v9.3.1, Oxford Instruments).

### Astrocyte enrichment and RNA isolation

At the prescribed interval, mice were anesthetized with 5.0% isoflurane before exsanguination and transcardial perfusion with ice-cold Dulbecco phosphate buffered saline (DPBS; Gibco # 14040133). Following perfusion, brain tissues were removed and quickly dissected to isolate the ipsilateral pericontusion cortex (or analogous region in sham mice) and dorsal hippocampal structure. Dissected tissues were immediately transferred into gentleMACS C-tube (Miltenyi #130-093-237), containing Adult Brain Dissociation Kit enzymatic digest reagents (Miltenyi #130-107-677), prepared according to manufacturer’s protocol. Tissues were dissociated using the “37C_ABDK” protocol on the gentleMACS Octo Dissociator instrument (Miltenyi #130-095-937) with heaters attached, following manufacturer’s suggested protocol. The resultant single cell suspension was used for magnetic bead enrichment for astrocytes (Miltenyi #130-097-678), following manufacturer’s suggested procedures, but utilizing three MS-columns (Miltenyi #130-042-201) to enhance purification. Washes were conducted using AstroMacs separation buffer (Miltenyi #130-117-336). The astrocyte cell surface antigen 2 positive fraction (ACSA-2^pos^) was collected into a separate tube, then centrifuged for 5 minutes at 1000x*g* at 4°C to pellet cells. Both the flow ACSA-2^neg^ and astrocyte enriched ACSA-2^pos^ fraction were collected from a subset of samples to validate putative gene enrichment efficiency.

Following centrifugation, supernatant was carefully aspirated, and cell pellets were lysed using RLT+ buffer containing 2-Mercaptoethanol using the Qiagen RNeasy+ Micro Kit (Qiagen #74034), following manufacturer’s suggested protocol for RNA isolation. RNA quantity was assessed using NanoDrop 2000 spectrophotometer, and approximately 25ng of total RNA was converted to cDNA using High-Capacity cDNA Reverse Transcription Kit (Applied Biosystems # 4368813). Resultant cDNA was stored at −80°C until assayed.

### Gene expression arrays

Multiplexed gene expression profiling was conducted on a ViiA7 qRT-PCR machine (Applied Biosystems) using a custom built Taqman low-density array card (Applied Biosystems) consisting of 44 genes of interest plus one housekeeping gene (*HPRT,* Mm00446968_m1). Genes on the array were curated from recent publications that previously determined putative astrocyte-specific responses to injury, disease, and aging [28, 29, 34]. Taqman gene probes were: *Amigo2* (Mm00662105_s1), *Apoe* (Mm01307193_g1), *B3gnt5* (Mm01952370_u1), *C1qa* (Mm00432142_m1), *C1qb* (Mm01179619_m1), *C3* (Mm01232779_m1), *CD109* (Mm00462151_m1), *CD44* (Mm01277161_m1), *Clcf1* (Mm01236492_m1), *Clu* (Mm01197002_m1), *CXCL10* (Mm00445235_m1), *Emp1* (Mm00515678_m1), *Fbln5* (Mm00488601_m1), *Fkbp5* (Mm00487406_m1), *Gbp2* (Mm00494576_g1), *GFAP* (Mm01253033_m1), *Ggta1* (Mm01333302_m1), *Gpc4* (Mm00515035_m1), *Gpc5* (Mm00615599_m1), *H2-*T23 (Mm00439246_g1), *Iigp1* (Mm00649928_s1), *Lcn2* (Mm01324470_m1), Maoa (Mm00558004_m1), *Osmr* (Mm01307326_m1), *Psmb8* (Mm00440207_m1), *Ptgs2* (Mm00478374_m1), *Ptx3* (Mm00477268_m1), *S100b* (Mm00485897_m1), *S1pr3* (Mm02620181_s1), *Serpina3n* (Mm00776439_m1), *Serping1* (Mm00437835_m1), *Sparcl1* (Mm00447784_m1), *Sphk1* (Mm00448841_g1), *Stat3* (Mm01219775_m1), *Steap4* (Mm00475405_m1), *Tgm1* (Mm00498375_m1), *Thbs1* (Mm00449032_g1), *Thbs2* (Mm01279240_m1), *Thbs4* (Mm03003598_s1), *Timp1* (Mm01341361_m1), *Tm4sf1* (Mm00447009_m1), and *Vim* (Mm01333430_m1). cDNA from each sample was diluted with TaqMan Gene Expression Master Mix (Applied Biosystems # 4369016), according to manufacturer’s protocol. Relative gene expression ratios were analyzed using the 2^-ΔΔCT^ method, with young-sham as the reference group. All gene expression ratios were Log_2_ transformed.

*Individual TaqMan assays*. Single assay reactions utilized for ACSA-2^pos^ sample enrichment calculations were also conducted on the ViiA7 using cDNA diluted with TaqMan Fast Advanced MasterMix (Applied Biosystems #4444557); *HPRT,* (Mm00446968_m1) *Aldh1l1* (Mm03048957_m1), *Tmem119* (Mm00525305_m1), *Dlg4* (Mm00492193_m1), *Klk6* (Mm00478322_m1), and *Nostrin* (Mm00724960_m1). Relative gene expression ratios were analyzed using the 2^-ΔΔCT^ method, with the ACSA-2^neg^ fraction as the reference group. All gene expression ratios were Log2 transformed.

### Statistical Analyses

All data were captured in a blinded manner, with the investigator unaware of groupings. The data codes were revealed and groupings assigned only after all data for each endpoint were captured. Statistical analyses were conducted in JMP Pro (v14.0), along with figures created in GraphPad Prism (v8.0). Pre-planned contrasts to examine the effect of age by post-surgical interval (histological and gene expression) were conducted using ANOVA with Sidak’s multiple comparison post hoc analysis. For heatmap of gene expression data, column (e.g. gene) data were standardized via z-score transformation, data were examined as a function of deviation from young sham values using ANOVA with Dunnet’s multiple comparison post hoc analyses. For representative analyses examining the effect of age within a post-injury interval, ANOVA with Sidak’s multiple comparison post hoc analysis was used. Multidimensional reduction was conducted using Principal Components Analysis (PCA) with Varimax orthogonal rotation for gene expression data. PCs were considered of interest using cutoffs of >1 for eigenvalue and Scree plot criteria. Resultant PC scores were calculated using regression method and their PC loadings were represented using arrows, where gauge (e.g. thickness) is indicative of loading magnitude, coloring heat of orange to blue indicates loading value with orange indicating greater magnitude and blue representing more minimal. Loading magnitudes with a cutoff of >|0.45| were graphically represented for interpretation. Significance for all measures was assessed at *p*<0.05.

## Results

### TBI elicits delayed reactive gliosis that is protracted in the aged brain

To examine whether reactive astrogliosis is sensitive to age at the time of injury, as well as the progression of time after injury, we examined GFAP reactivity in the ipsilateral dorsal hippocampal formation. GFAP, an intermediate filament protein along with Vimentin, are classical histological markers for assessing reactive astrogliosis, which are consistently assessed to gauge trauma-associated responses of astrocytes [35–37].

Further, utilizing these markers is sufficient to assess the degree of homeostatic disturbance, as GFAP and Vimentin have shown a somewhat linear relationship with severity of- and proximity to injury in the brain [25]. We found a TBI-induced increase in hippocampal (HPC) GFAP^+^ astrogliosis over the first 7 days post-injury in both young and aged animals (**Figure 1A**/B). Young mice showed the highest magnitude response by 3 days post-injury and a return toward baseline thereafter, whereas the aged cohort exhibit a protracted response with the highest magnitude response at 7 days post-injury (**Figure 1A**/B). Similarly, we quantified Vimentin in the context of another reactive marker associated with astrocytes: S100B. Although S100B is predominantly expressed by astrocytes in the CNS, it is not exclusive to this cell type [38], therefore examining the co-labeled fraction with Vimentin allowed us to assess its astrocyte-specific reactivity. Examining the colocalized area (e.g. dual positive for Vimentin and S100B), our data demonstrated that in the young brain, these dual-positive astrocytes are less reactive to TBI as assessed by positive area (**Figure 2A**/B), compared to the aged brain at all post-injury intervals.

**Figure 1.**
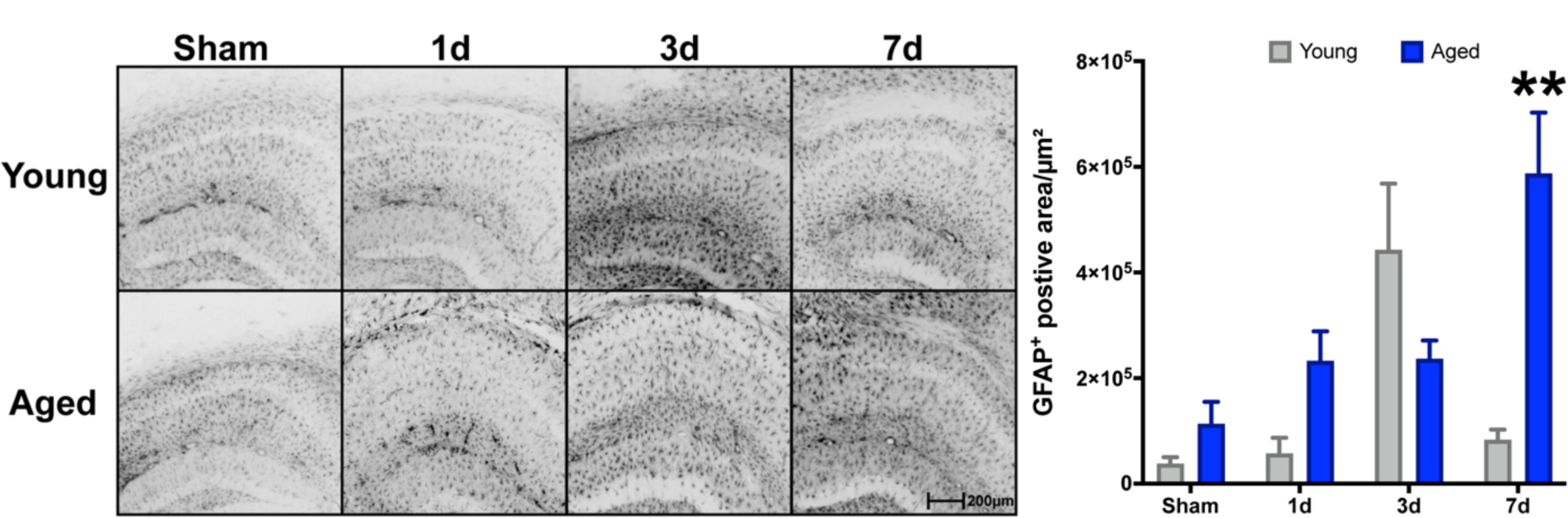
Age-related progressive reactivity of GFAP^+^ astrocytes in the hippocampus after TBI. Young (4-mo) and aged (18-mo) mice were subjected to sham or CCI injury, and euthanized at 1, 3, or 7 days post-injury. Serial sections comprising the dorsal hippocampus were imaged for GFAP labeling and quantified as the pixel positive area for each of the 8 groups. GFAP^+^ threshold was set on young sham tissue. TBI induced a progressive increase of GFAP^+^ astrocyte staining in aged mice (n=5/group) through 7 days post-injury, whereas the GFAP^+^ staining intensity in young mice (n=4/group) was most prominent by 3 days post-injury. Data were analyzed using two-way ANOVA with Sidak’s post hoc correction examining pairwise interactions for each time interval. ANOVA revealed significant differences due to age (F (1, 28) = 7.924, P=0.0088), interval (F (3, 28) = 7.593, P=0.0007), as well as their interaction (F (3, 28) = 9.065, P=0.0002). \*\**p*<0.0001, for pairwise comparison of 7d interval between young and aged. Data are presented as mean±SEM. Young = gray bars, Aged = blue bars. Scale bar is 200μm.

**Figure 2.**
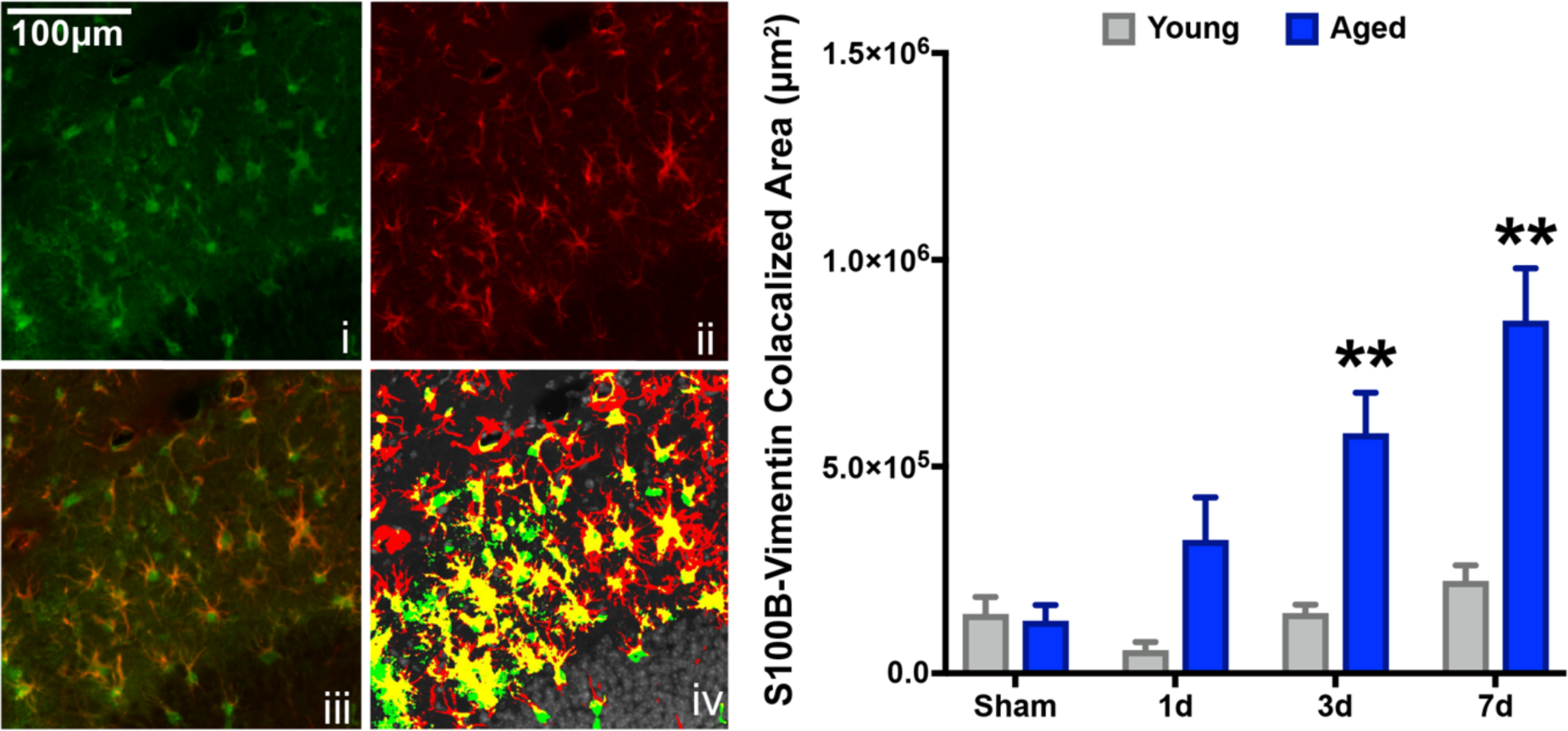
Aged mice display exacerbated S100B^+^Vim^+^ reactive astrogliosis in the hippocampus following TBI. Serial sections comprising the dorsal hippocampus were imaged for S100B (**i**) and Vimentin (**ii**) as a colocalization image (**iii**). Pixel positive area was defined for each stain using young sham’s levels. HALO colocalization algorithm was used to compute the total colocalized area for both S100B (green, **iv**) and Vimentin (red, **iv**), which was represented as the dual-positive staining fraction (yellow, **iv**). Dual-positive astrocytes were increasingly reactive as a function of time after injury for aged mice, which show a progressive accumulation peaking at 7 days post-injury. Comparatively, young mice display relatively little change as a result of TBI at any time post-injury. ANOVA revealed significant differences due to age (F (1, 28) = 33.61, P<0.0001), interval (F (3, 28) = 10.38, P<0.0001), and their interaction (F (3, 28) = 5.831, P=0.0032). \*\**p*<0.01 for pairwise comparisons of 3d and 7d interval between young and aged mice. n=4-5/group. Data are presented as mean±SEM. Young = gray bars, Aged = blue bars. Scale bar is 50μm.

### Advanced age promotes accumulation of clasmatodendrosis in astrocytes in the HPC

First reported by Alois Alzheimer, and later coined by Cajal [39], clasmatodendrosis is typified by the beading and diminishment of astrocyte projections, paired with vacuolization and swelling of the cytoplasm [40–42]. This astrocyte pathology has been observed in AD, Binswanger’s mixed dementia, aging, and ischemic brain tissues [41–45]. Furthermore, recent evidence in post-mortem analyses of Chronic Traumatic Encephalopathy (CTE) has demonstrated trauma-induced accumulation of a morphologically distinct subset of reactive astrocytes [46]. Characteristically, these astrocytes had a beaded ‘pearl on a string’ degenerative morphology. We observed a significant accumulation of these degenerative-like astrocyte morphologies in the molecular layer of the stratum radiatum of the hippocampus (**Figure 3A**) in our injured aged mice, and these astrocytes were found in aggregates (**Figure 3B**). Our triple labeling method revealed S100B^+^Vimentin^+^ dual-positive vacuolization along GFAP^+^ filament tracks and somatic hypertrophy (**Figure 3C**) consistent with morphological hallmarks attributed to clasmatodendrosis of astrocytes. Quantification of the numbers of astrocytes exhibiting clasmatodendrosis showed that only aged mice showed these morphologically distinct astrocytes in the HPC, and that TBI in the aged mice induced a decrease in these astrocytes over the 3-day post-injury period, which returned to pre-TBI levels by 7 days post-injury (**Figure 3A**).

**Figure 3.**
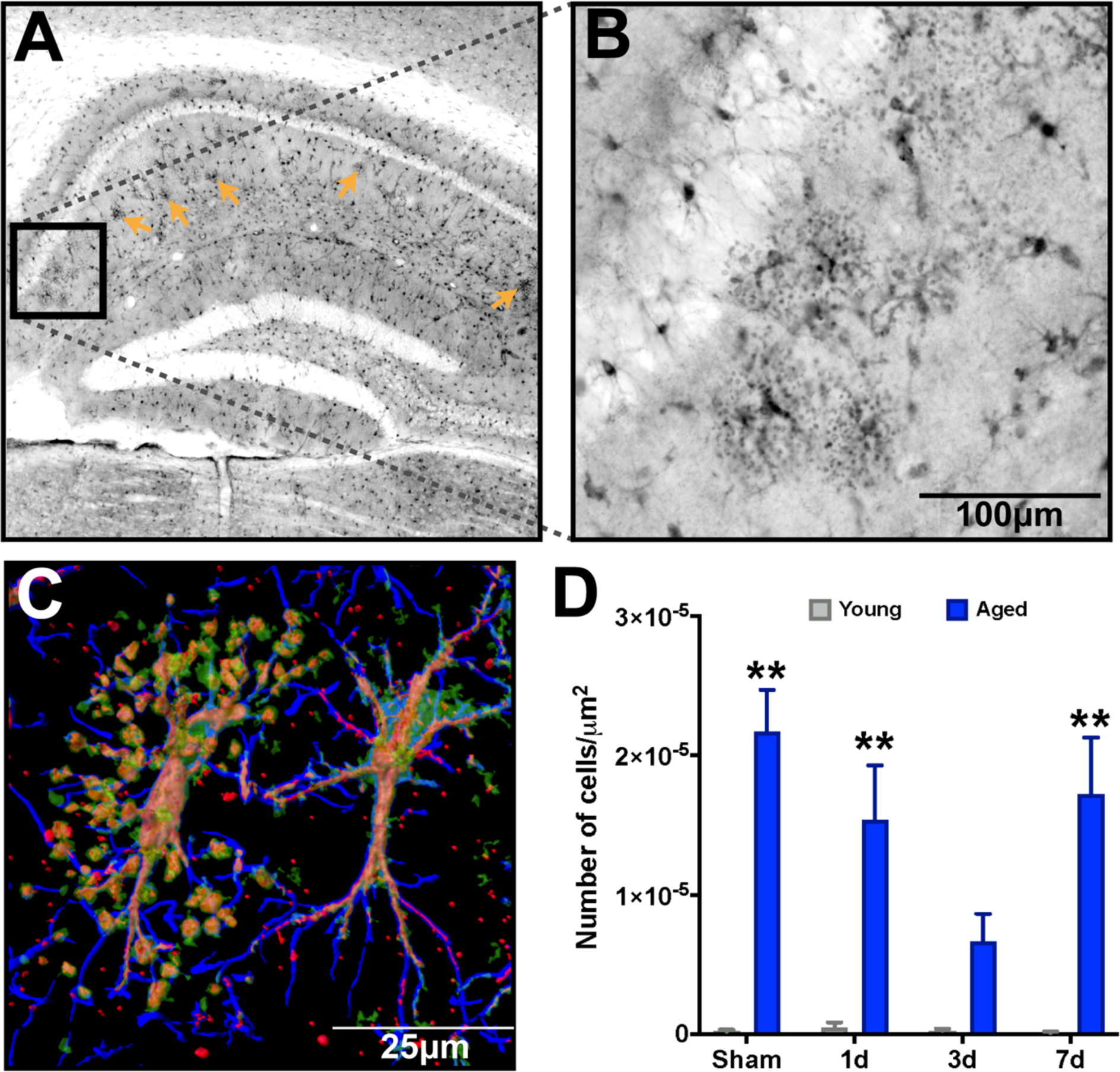
Advanced age results in accumulation of clasmatodendrosis in the stratum radiatum of the dorsal hippocampus. S100B labeling reveals a significant accumulation of degenerative astrocytes **A.** Low power magnification of dorsal hippocampus shows the distribution of clasmatodendrosis localized within the stratum radiatum of the CA1 region of the hippocampus (orange arrows). **B.** High power magnification shows the distinctive enlarged soma and vacuolization of processes distinctive to clasmatodendrosis for a representative cluster of astrocytes. **C.** Imaris surface render of a confocal z-stack of GFAP (blue), S100B (green), and Vimentin (Red) demonstrates an astrocyte with clasmatodendrosis (left) showing the co-localization of S100B^+^Vimentin^+^ beads along GFAP^+^ processes, and a non-degenerative morphology reactive astrocyte adjacent to it (right). **D.** There is a significant accumulation of clasmatodendrosis in the aged brain, which did show some temporal response to TBI at 3 days post-injury; however by 7 days, levels return to approximate those in the uninjured condition. n=4-5/group. Data were analyzed using two-way ANOVA with Sidak-Holm post hoc correction examining pairwise interactions for each time interval. ANOVA revealed a significant main effect due to age (F (1, 28) = 64.41, P<0.0001), however neither interval (F (3, 28) = 2.800, P=0.0590), nor their interaction were significantly different (F (3, 28) = 2.856, P=0.0557). \*\**p*<0.01 for pairwise comparisons of sham, 1d, and 7d interval between young and aged mice. n=4-5/group. Data are presented as mean±SEM. Young = gray bars, Aged = blue bars.

### Astrocyte endfoot integrity is diminished by TBI and advanced age

Consistent evidence has implicated astrocytes as a critical component of the inflammatory response to CNS trauma [47], mediated in part by their polarization of endfeet around vascular surfaces throughout the brain. Classically, the polarity of astrocytes to perivascular localization has been assessed by analysis of aquaporin-4 (Aqp4), which is a water channel expressed exclusively in the CNS by astrocytes [48] and is a critical component of waste clearance pathways. Given these critical physiological properties, we quantified the total area of Aqp4 staining in the ipsilateral HPC of our young and aged cohorts. Our data demonstrate that there is a trend for aged mice (sham) to have decreased Aqp4 positive area, relative to young shams (**Figure 4**). However, after TBI, we observed a significant change in Aqp4 only in the aged mice, and only at 3 days post-injury (**Figure 4**). The TBI-induced rarefaction of Aqp4^+^ seen at 3 days post-injury returned to similar levels of their young comparators by 7 days post-injury.

**Figure 4.**
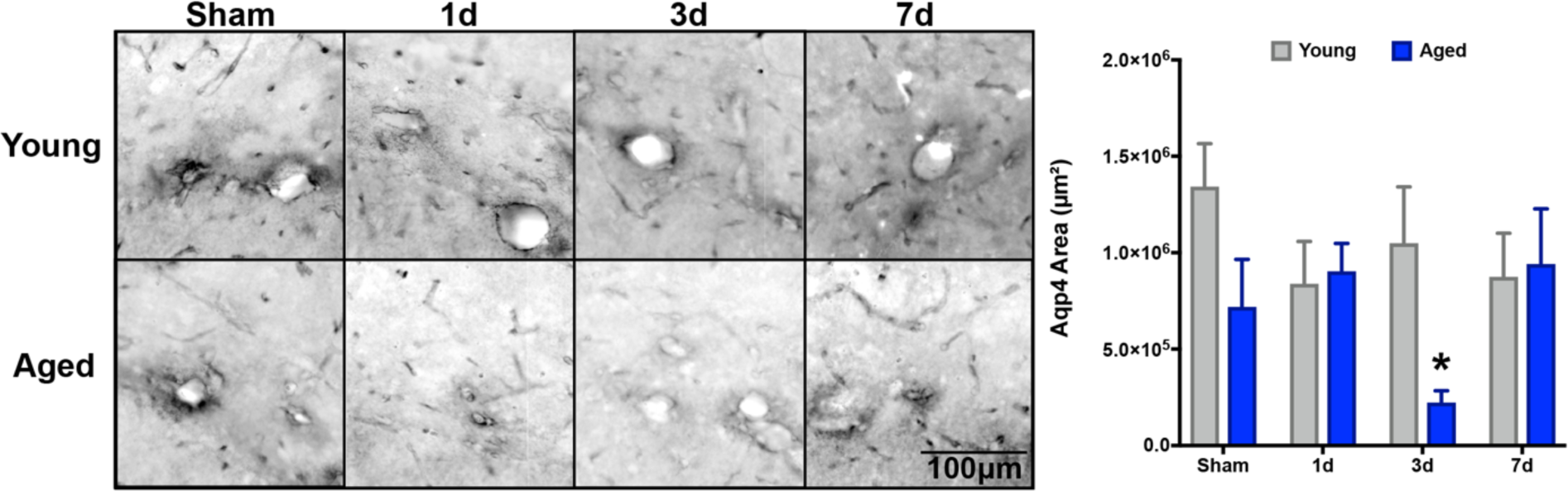
Aquaporin-4 displayed a delayed transient response to TBI in the aged brain. Serial sections comprising the dorsal hippocampus were imaged for aquaporin-4 (Aqp4) labeling and quantified as the pixel positive area for each of the 8 groups. Aqp4^+^ threshold was set on young sham tissue. Overall, there was a significant decrease in Apq4 staining due to advanced age (F (1, 28) = 4.506, P=0.0435), which was significantly reduced compared to young at the 3d post-injury interval (*p<0.05). This age-related loss of Aqp4^+^ recovered to sham-like levels at the 7d post-injury interval. There were no significant overall differences observed for due to interval (F (3, 28) = 1.144, P=0.35), nor an interaction effect (F (3, 28) = 2.292, P=0.10). Data were analyzed using two-way ANOVA with Sidak-Holm post hoc correction examining pairwise interactions for each interval. n=4-5/group. Data are presented as mean±SEM. Young = gray bars, Aged = blue bars.

### Profiling TBI-induced astrocyte transcriptional responses across representative A1/A2 categorizations

Recent work has defined astrocyte-specific responses to either generic, microglia/lipopolysaccharide (LPS) mediated, or middle cerebral artery occlusion (MCAO) stimuli in neonate rodents to fall within categorical bins of ‘Pan reactive’, ‘A1’, or ‘A2’, respectively [34, 49]. Given that TBI elicits upregulation and reactivity of a variety of convergent pathways attributed to both microglial and ischemic responses, we examined a subset of these transcriptional markers that were prominent in defining these categorizations, with an additional focus on genes associated with regulating synaptic function [29]. Therefore, we wanted to determine whether TBI and/or age affected transcriptional responses of astrocytes. In order to examine putative astrocyte responses in the context of age, injury, and interval, we utilized a magnetic bead enrichment protocol (**Figure 5A**), previously demonstrated by others to enrich astrocytes from adult CNS tissues [50–52]. Using this procedure, we validated the enrichment efficiency by examining genes enriched for putative CNS subsets of astrocytes (*Aldh1l1)*, neurons (*Dlg4*), oligodendroglia (*Klk6*), microglia (*Tmem119*), and endothelia (*Nostrin*). The results (**Figure 5B**) demonstrated marked enrichment of *Aldh1l1* in the ACSA-2^+^ fraction relative to the markers for other cell types.

**Figure 5.**
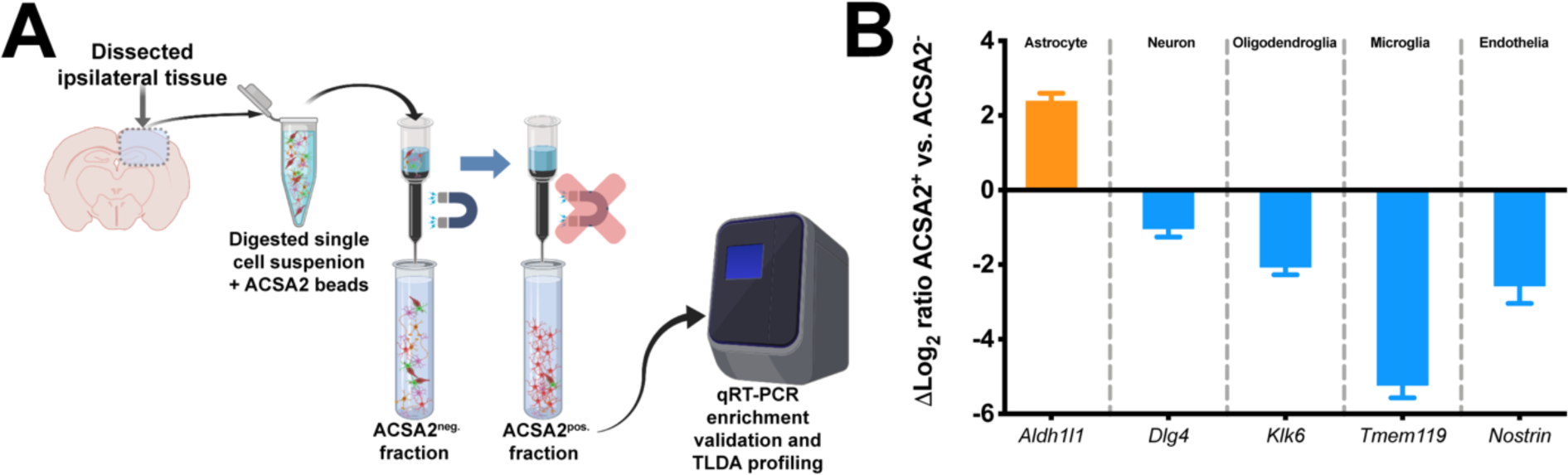
Validation of ACSA-2 astrocyte cell enrichment from 4-month-old mouse brain tissue. **A** Generalized workflow for ACSA-2 magnetic bead enrichment of astrocytes from the injured brain parenchyma. Digested cell suspensions were labeled with the ACSA-2 magnetic bead before being placed in the magnetic column for removal of non-specific cells, which were collected into a tube as the ACSA-2^neg.^ fraction. Removal of the column from the magnetic stand allowed the flow-through of the retained ACSA-2 astrocytes to be collected as the ACSA-2^pos.^ fraction. RNA from both fractions was harvested to examine gene expression endpoints. **B.** Gene expression analyses using putative markers of five neural tissue subsets; astrocyte (*Aldh1l1)*, neuronal (*Dlg4*), oligodendroglial (*Klk6*), microglial (*Tmem119*), and endothelial (*Nostrin).* These data demonstrate a significant induction of putative astrocyte signature (orange bar), with little signature of other cell populations (blue bars). Log_2_ fold change is a ratio of ACSA-2^pos^ to ACSA-2^neg^. TLDA =Taqman low density array cards.

Using RNA from magnetic bead enriched astrocyte fractions, we profiled 44 genes that spanned 4 categorical bins attributed to astrocyte biological and pathological profiles (**Figure 6A**). Overall, our findings indicate that neither TBI nor advanced age were particularly constrained to either A1 or A2 categorical bins, with significant gene expression changes observed predominantly in the ‘pan-reactive’ and ‘synaptic modifying’ motifs (**Figure 6A**; black and magenta headers). However, we did observe the induction of several genes in the ‘MCAO-associated’ bin (**Figure 6A**; green header). Notably, there are minimal transcriptional changes attributed to TBI or advanced age in the reactive microglia/LPS-associated ‘A1’-associated phenotype (**Figure 6A**; blue header). Although neither TBI nor age were found to predominantly align with these categorizations, we did observe several interesting dynamics when examining the effect of age at each temporal post-surgical interval. For *CD44, CXCL10, C1qa, Amigo2, Ligp1, and B3gnt5* we observed significant differences occurring at 1 day post-injury for aged astrocytes, compared to young (**Figure 6B**). Notably, there seems to be a plateau effect with these markers initiated at 1 day that had a protracted response in magnitude trends carrying through 7 days post-injury (**Figure 6B**). Comparatively, both *GPC6* and *Cd109* were consistently depressed in expression in aged mice relative to their young counterpart at all time intervals and showed relatively little fluctuation as a function of post-injury interval (**Figure 6B**).

**Figure 6.**
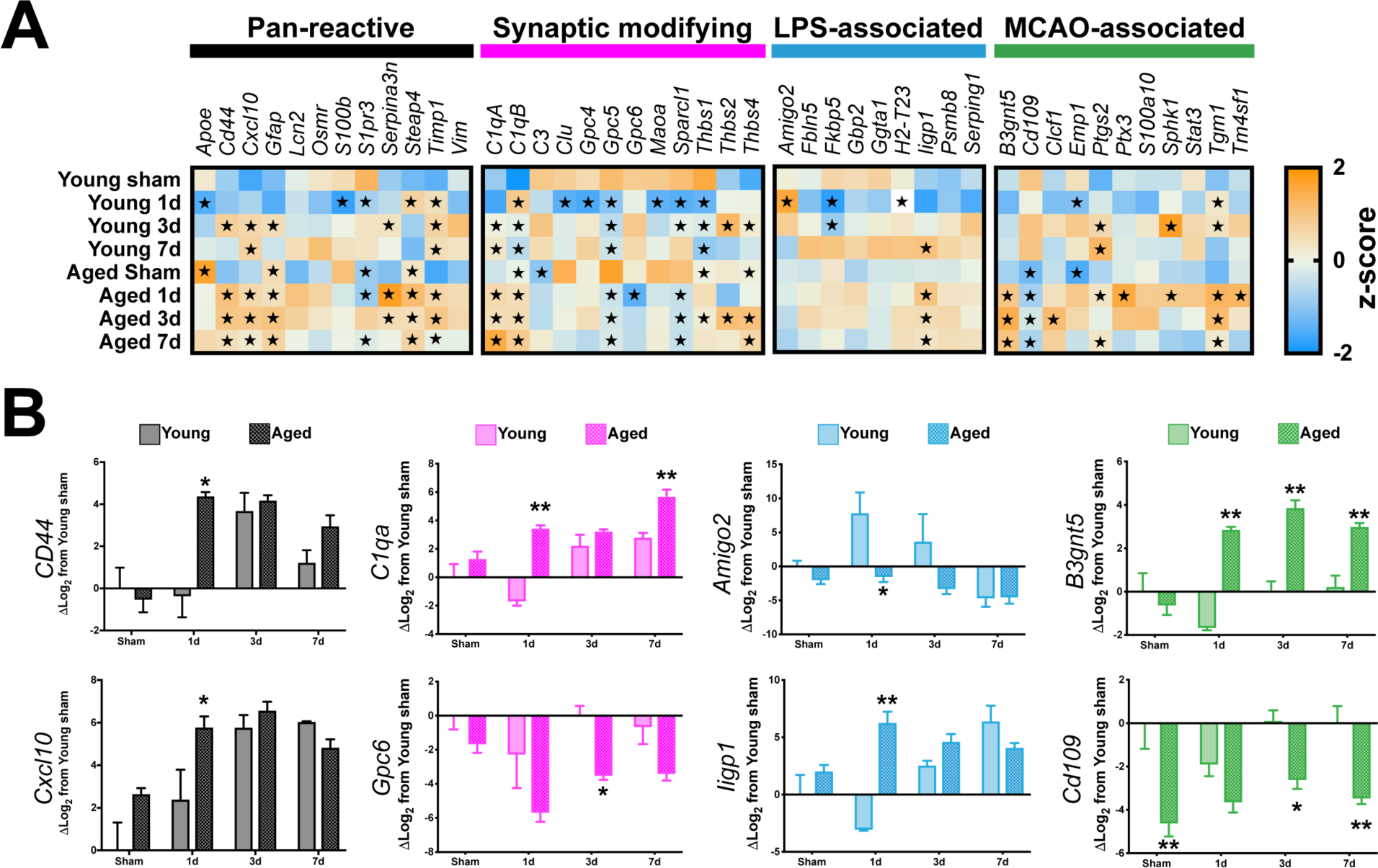
Examining TBI and age-related astrocyte-specific responses. **A** Gene expression responses for the 8 groups were quantified via custom TaqMan low-density array (TLDA) cards, comprising 44 genes of interest that spanned previously defined categorical bins attributed to ‘Pan-reactive’ (black bar), ‘Synaptic modifying’ (magenta bar), ‘LPS-associated’ (“A1”, blue bar), or ‘MCAO-associated’ (“A2”, green bar) responses of astrocytes. Neither TBI nor advanced age were exclusively confined to either of the A1/A2 phenotypes, whereas the bulk of significant deviations from young-sham were found within the ‘Pan-reactive’ or ‘Synaptic modifying’ gene sets. Each square shows the mean Z-score of Log_2_ expression as a gene/group representation. **B.** Representative subset of genes for each category demonstrates age-related pair-wise comparisons. Gene expression data are relative to young sham values. All data were Log_2_ transformed. For panel A, data were analyzed using a two-way ANOVA with Dunnet’s post hoc correction, with young sham defined as the ‘control’ value. Data are presented as the mean Z-score for each gene, with orange representing positivity, and blue representing negative. ★ denotes *p*<0.05 relative to young sham. For panel B, data were analyzed using two-way ANOVA with Sidak-Holm correct to examine pair-wise age-related interactions with \**p*<0.05 or *\*\*p*<0.01 indicating significant effects due to age within the pairwise comparison interval. n=4-5/group. Data are presented as the mean±SEM.

### Examining astrocyte-specific transcriptional dynamics in an unbiased multivariate approach

We have previously examined the brain’s response to TBI focusing upon myeloid (e.g. microglia and monocytes) specific responses utilizing multivariate methods to delineate the effect of injury and time [32]. Herein, we have employed similar methods using principal components analyses (PCA) in an unbiased approach to examine how age, interval, and injury converge to drive the multidimensional responses of astrocytes in the injured milieu. This analysis yielded three orthogonal PCs, in total accounting for 64.3% of the total variance in astrocyte transcriptional responses within the 46 genes examined (**Figure 7A**). The PC1 module reflects gene expression shifts due to TBI as 1,3, and 7 day groups that were largely distinct from their sham comparators. Interestingly, there is an initial divergence in this profile as a function of age (**Figure 7B**) at the 1 day interval, however by 7 days PC loadings between age groups were strikingly similar. The PC1 module was largely driven by *CD44, Serpina3n,* and *Tgm1* gene expression profiles (**Figure 7C**; PC1 loadings >0.8). Comparatively, the PC2 module reflects the temporal dynamics of the post-injury intervals, where both young and aged astrocytes display similar temporal u-shaped trajectories, with an initial repressed response, which overshoots its baseline for young, but fails to return for aged (**Figure 7B**). PC2 responses were largely driven by expression changes associated with *Apoe, Maoa, Clu, S100b, Gpc5, and Sparcl1* (**Figure 7C**; PC2 loadings >0.8). Lastly, the PC3 module encapsulates gene expression profiles predominated by advanced age, as the sham, 1 day, 3 day, and 7 day aged astrocytes were starkly delineated in the multivariate plot (**Figure 7A**/B), showing no overlap at any point, comparatively. Aged responses driving the PC3 module were largely attributed to *CD109* (**Figure 7C**; PC3 loadings >0.8). Lastly, we examined the similarity between our dataset and a previously published RNAseq dataset by Anderson *et al*. (2016) that examined astrocyte-specific responses following spinal cord injury (SCI). We chose to examine the 3 day post-injury interval of our gene expression responses, because this time point showed several convergent responses across the endpoints described above. Using the mean log transformed fold change (logFC) from ‘SCI WT astrocytes’ vs. ‘Uninjured WT astrocytes’ from the Anderson *et al*. dataset, we plotted the mean logFC of our genes that had a PC loading >|.45| from our 3 PCs for both young and aged astrocytes at the 3 day post-injury interval (**Figure 7D**).

Comparatively, our data follows remarkably similar expression profiles, particularly for PC1. Preservation of these trends between our current findings and those in the Anderson *et al*. (2016) dataset may potentially suggest a conservation of trauma-induced transcriptional profiles in astrocytes irrespective of CNS locale.

**Figure 7.**
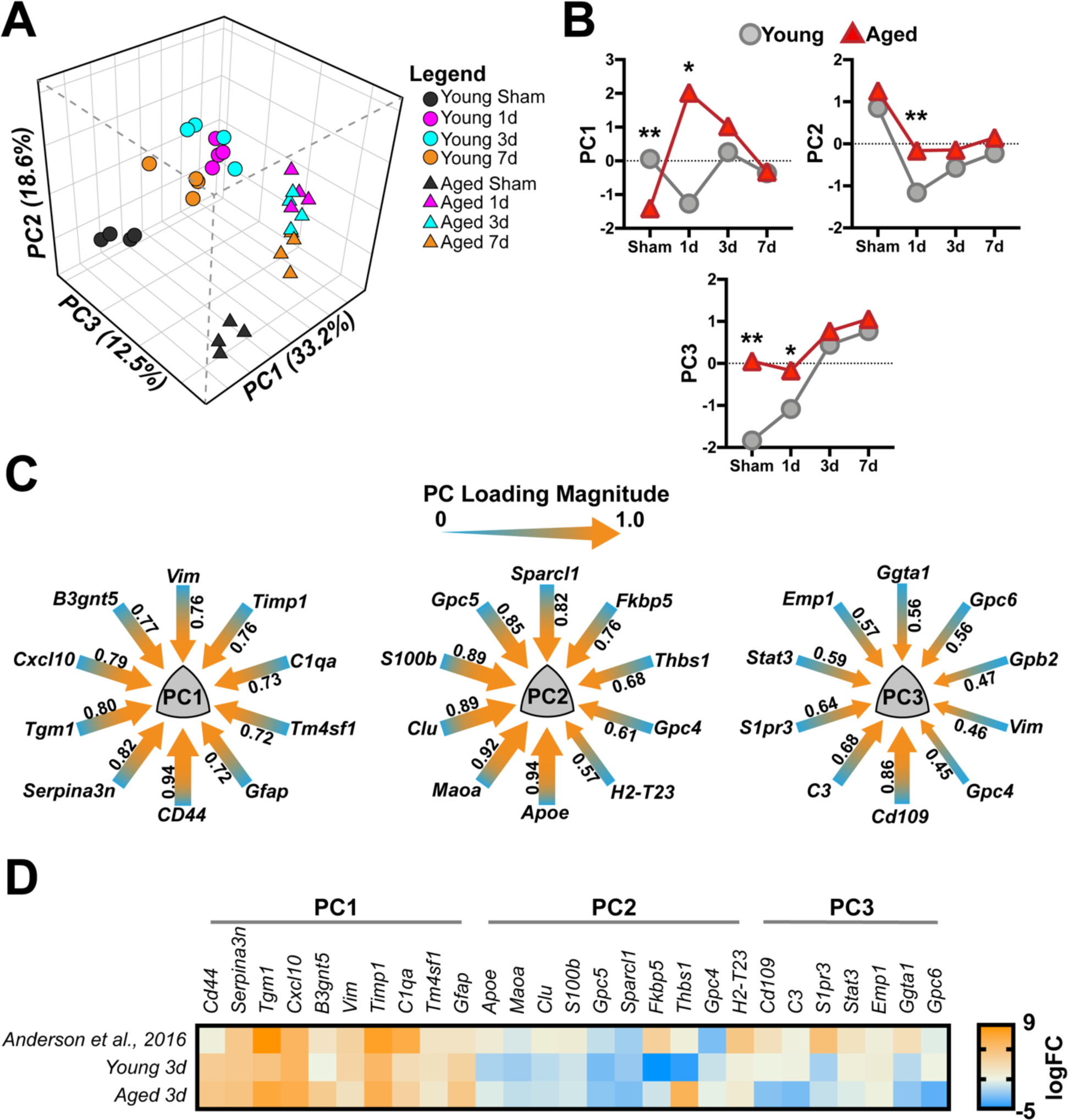
Astrocyte-specific response patterns to age and TBI revealed by principal components analysis (PCA) **A.** Multivariate dimensional reduction by PCA uncovered 3 orthogonal PC groups, that cumulatively accounted for 64.3% of the total variance. Within this multidimensional space, PC1 (33.2% variance) reflected a primarily TBI-induced response, while PC2 (18.6% variance) reflected some of the temporal responses within the dataset, and lastly PC3 (12.5% variance) reflected an age-related effect. **B.** Temporal trajectories of the mean PC score for each group in each of the three PCs identified. Data were analyzed using two-ANOVA, with Sidak-Holm correction for examining pairwise interactions for each time interval. Data points represent the mean PC score for each interval and age. Young = gray circles, Aged = Red triangles. **C.** PC loading magnitudes for the top 10 genes >|.45| loading for each of the three PCs. Arrow gauge (thickness) is proportional to loading magnitude, with heat (color) representing loading directionality toward 1.0. **D.** Comparison of the Log_2_ fold change (logFC) from “SCI WT Astrocyte” vs. “Uninjured WT Astrocytes” from Anderson *et al*., 2016 versus mean logFC of young 3 day and aged 3-day cohorts from the current study. Comparing 3-day TBI responses from both young and aged cohorts with SCI-induced responses reveals a striking similarity in PC1-related profiles between the two datasets. Data were arranged within the previously defined PCs from above, duplicates of genes from PCs (e.g. PC1 *Vim*) were removed for clarity. Additionally, no data were found in the Anderson *et al*. dataset for *Gpb2*, which was also removed for clarity.

## Discussion

Our study demonstrates that TBI induces a progressive increase of reactive astrogliosis in the aged hippocampus. We demonstrated novel heterogeneity between reactive astrocytes’ morphologies and age-related clasmatodendrosis of astrocytes. Further, after dissecting the molecular phenotypes using a focused array, our current data demonstrate that TBI was not sufficient to align itself to previously defined molecular profiles associated with the “A1/A2” phenotypic bins [28, 34]. However, using these unique gene expression markers did identify several novel responses of aged astrocytes to TBI. Unbiased multivariate analyses demonstrated that injury, interval, and age correlated with distinct gene expression profiles collectively pointing toward increased inflammatory response, decreased synaptic support, and increased pruning phenotypes of aged astrocytes following TBI, compared to young. These findings suggest that astrocytes in the aged brain have lost key restraining- or acquired dysfunctional mechanisms, ultimately driving what we hypothesize to be a maladaptive response to TBI, compared to young.

In terms of the age-related progressive reactive astrocyte response in the hippocampus, our findings parallel a previous report that examined GFAP reactivity in 21- to 24-month-old mice that received a CCI centered over the caudate putamen [53]. Further, although our data demonstrate an exacerbation of the S100B response in the aged brain, the exact consequence of overactive S100B in the aged brain remains unknown, as it has been implicated in a variety of roles for both healthy and diseased CNS [54]. Whether aged astrocytes are propagating damage-associated molecular patterns (DAMPs) via S100B [55], or this signaling is potentiating previously defined neurosupportive roles [56–59] remains to be determined. Given the exacerbated responses of both of these measures, they may lend themselves particularly well as prognostic indicators for aged individuals acutely after TBI [60]

Our present findings demonstrating an age-related increase in clasmatodendrosis are corroborated by previous work examining human brain [43]. We observed the aggregation of this morphologically distinct subset of astrocytes confined to the stratum radiatum of the CA1, which recapitulates previous work in rodents demonstrating this as a susceptible region for accumulating these astrogliopathies [61–63]. Although the exact role or cause of this pathology is not well-defined, recent reports have demonstrated corollaries with senescence, autophagy, metabolic dysfunction, ER stress, and NFκB signaling [61, 62, 64, 65]. Moreover, our findings potentially highlight a trauma-induced turnover response occurring at the 3 day post-injury interval, such that the age-related accumulation of these degenerative astrocytes may be susceptible to inflammatory-mediated removal or clearance mechanisms, which remains to be elucidated.

Aqp4 expression and distribution across the perivascular environment has been previously shown to be significantly disrupted as a function of aging alone in a rodent model of aging and in human post-mortem examinations [66, 67]. Critically, Aqp4 serves as an integral water channel in the brain’s glymphatic system for waste material clearance by facilitating the exchange between both CSF and interstitial fluids [68]. These dynamics are largely associated with the concentrated Aqp4 localization along the perivascular and subpial astrocyte endfoot membranes [69]. Our current findings recapitulate previous work demonstrating an age-related reduction in Aqp4 staining intensity [66, 67]. Moreover, the transient responses we observed in response to TBI may point toward a refractory period wherein astrocytes have lost their vascular-related homeostatic capacity in the aged brain, potentially implicating transient impairment of solute clearance mechanisms, vascular tone, as well as cerebral blood flow.

Recent work profiling the transcriptome of aged astrocytes has demonstrated altered expression in pathways associated with synaptic function, inflammatory disposition, and disproportionate response to LPS-mediated insult [28, 29]. In essence, this previous work implicates aged astrocytes as a “primed” cell population, similar to how microglia have been shown to become predisposed to insult or stress in the aged brain [70]. At least in the context of our narrow geneset, our current findings would be in agreement with the potential for TBI to initiate a phenotype outside these bounds, or as a continuum of these stimulus-specific bins [49]. Several unique responses that included age-related shifts in expression for inflammatory markers *CXCL10, CD44,* and *C1qa*. Our data demonstrate a similar exacerbated CXCL10 response in aged astrocytes due to TBI. CXCL10 has been shown to modulate chemotaxis and initiate cross-talk between immunoreactive microglia and astrocytes [71], and was previously documented to have an exacerbated response in aged astrocytes following LPS-induced inflammatory challenge [28]. In this sense, CXCL10 in the aged brain could be potentiating aberrant cross-talk with microglia. CD44 defines astrocyte precursors [72, 73] their tissue heterogeneity in the adult brain [74], as well as being both injury and disease responsive [75–77]. Interestingly, recent work would suggest that the acquisition of CD44 on astrocytes is indicative of a regressive phenotype toward an immature state [75]. Collectively, our findings of increased inflammatory response paired with reductions in synaptic support markers, such as Gpc6 and the upregulation of C1qa, a complement protein that has been shown to play a role in tagging synapses for their selective removal [11], potentially suggest a maladaptive environment where neurons and/or their synapses are highly susceptible to detrimental phenotypes by aged astrocytes.

Our multivariate approach to interpreting these responses yielded several uniquely correlating gene expression phenotypes, which show that in both the uninjured and 1 day post-injury timepoints there were significant dysfunctional outcomes within these multivariate domains, compared to young. Comparatively, when we examined our injury-induced responses that displayed some exacerbation in the context of age (e.g. PC1) with transcriptome profiling of injured astrocytes from the spinal cord, there was an analogous resemblance to these data [20]. Although injury was produced in the spinal cord, there are several analogous parameters with Anderson *el al*. (2016) with respect to the contusion injury and our CCI model. Therefore, concordance between these findings potentially point toward a conserved reactivity among astrocytes following contusion, irrespective of CNS locale.

Despite concordance between our findings with several previous reports for aging and injury, there remain several notable caveats with our design and resultant dataset that warrant acknowledgement as well as open exciting future directions to explore. Notably, our antibody-based enrichment method may induce bias in the types of astrocytes, only ACSA-2^pos^ enriched from our samples, compared to recent methods using a global astrocyte-specific (e.g. *Aldh1l1*) promoter to drive mechanisms for polyribosomal RNA purification (e.g. bacTRAP or RiboTag), as was previously described for profiling age-related responses for astrocytes [28, 29]. Similarly, there is known heterogeneity in astrocyte populations in homeostasis [23] as well as a function of distance from injury (for review see [26]). Therefore, it is possible that even though we observed significant differences in our aged cohort across several timepoints, there may be even more nuanced, but disease-relevant responses that would be detected using either of the ribosomal affinity purification techniques or single cell analyses in tandem with whole transcriptome profiling. Lastly, our focus in this study was to understand the acute responses of astrocytes, as a complement to our previous work showing altered neuroinflammatory responses due to injury and/or advanced age chronically. Future critical work examining the persistence of chronic dysfunctional responses in aged astrocytes may shed light on how these cells are affecting unique signaling domains underlying chronic neuronal, synaptic, and cognitive impairments seen in our TBI model [31, 32].

## Conclusion

Our results demonstrate that advanced age predisposes astrocytes to acquire degenerative physiologic properties, with progressively aberrant morphological responses to TBI, compared to young. In terms of the heterogeneous molecular responses attributed to astrocytes, our findings parallel previous reports in terms of their age-related altered basal signature, which is exacerbated by TBI. CCI-induced molecular signatures show a surprising conservation of phenotypes when compared with SCI. Taken together, our initial characterization of aged astrocytes’ response to TBI begs the question as to whether these disproportionate motifs are responsible for exaggerated functional outcomes in the aged brain, and importantly, whether targeting astrocytes specifically to alter some of these responses would offer functional recovery.

## Abbreviations

AD: Alzheimer’s disease
Aqp4: Aquaporin4
ACSA-2: Astrocyte Cell Surface Antigen 2
BBB: Blood brain barrier
CBF: Cerebral blood flow
CSF: Cerebral spinal fluid
CTE: Chronic Traumatic Encephalopathy
CCI: Controlled cortical impact
CA1: Cornu Ammonis area 1
DAMPs: Damage-associated molecular patterns
GFAP: Glial fibrillary acid protein
HPC: Hippocampus
LPS: Lipopolysaccharide
MCAO: Middle cerebral artery occlusion
NGS: Normal goat serum
PFA: Paraformaldehyde
PBS: Phosphate buffered saline
PCA: Principal Component Analysis
SCI: Spinal cord injury
TLDA: Taqman low density array
TBI: Traumatic brain injury

## Declarations

### Ethics approval and consent to participate

All procedures involving animals for this study were approved by the Institutional Animal Care and Use Committee of the University of Kentucky.

### Consent for publication

Not applicable.

### Availability of data and materials

The datasets generated and/or analyzed during this study are available from the corresponding author on reasonable request.

### Author contributions

ANE and JMM designed the research studies. ANE, AAG, ADB, and JMM performed the experiments. LJVE contributed critical reagents and provided manuscript edits. ANE, ADB and JMM analyzed the data, ANE and JMM wrote the manuscript. All authors have read and approved the final manuscript.

## Acknowledgements

We also wish to thank Edgardo Dimayuga for the excellent technical assistance toward the completion of this project.

